# Morphogenesis and dynamics of slime molds in various environments

**DOI:** 10.1101/622662

**Authors:** Fernando Patino-Ramirez, Aurèle Boussard, Chloé Arson, Audrey Dussutour

**Author notes:** Corresponding authors (FP), (AD). These authors contributed equally to this work (Chloé Arson, Audrey Dussutour).

## Abstract

Cells, including unicellulars, are highly sensitive to external constraints from their environment. Amoeboid cells change their cell shape during locomotion and in response to external stimuli. Physarum polycephalum is a large multinucleated amoeboid cell that extends and develops pseudopods. In this paper, changes in cell behavior and shape were measured during the exploration of homogenous and non-homogenous environments that presented neutral, and nutritive and/or adverse substances. In the first place, we developed a fully automated image analysis method to measure quantitatively changes in both migration and shape. Then we measured various metrics that describe the area covered, the exploration dynamics, the migration rate and the slime mold shape. Our results show that: 1) Not only the nature, but also the spatial distribution of chemical substances affect the exploration behavior of slime molds; 2) Nutritive and adverse substances both slow down the exploration and prevent the formation of pseudopods; and 3) Slime mold placed in an adverse environment preferentially occupies previously explored areas rather than unexplored areas using mucus secretion as a buffer. Our results also show that slime molds migrate at a rate governed by the substrate up until they get within a critical distance to chemical substances.

**Author summary:** *Physarum polycephalum*, also called slime mold, is a giant single-celled organism that can grow to cover several square meters, forming search fronts that are connected to a system of intersecting veins. An original experimental protocol allowed tracking the shape of slime mold placed in homogenous substrates containing an attractant (glucose) or a repellent (salt), or inhomogeneous substrates that contained an attractive spot (glucose), an eccentric slime mold and a repulsive spot (salt) in between. For the first time, the rate of exploration of unexplored areas (primary growth) and the rate of extension in previously explored areas (secondary growth) were rigorously measured, by means of a sophisticated image analysis program. This paper shows that the chemical composition of the substrate has more influence on the morphology and growth dynamics of slime mold than that of concentrated spots of chemicals. It was also found that on a repulsive substrate, slime mold exhibits a bias towards secondary growth, which suggests that the mucus produced during slime mold migration acts as a protective shell in adverse environments.

## Introduction

Large-scale spatial patterns in biology are common and knowing how these patterns evolve and what are their functional role, enables us to understand the evolution of biocomplexity (see e.g. (1–4)). Morphogenesis has been studied in length at the cell level (see e.g (5–8)); cells are highly sensitive to geometrical and mechanical constraints from their microenvironment and respond to these conditions by changing shape (see e.g. (9,10)); these transformations impact cell migration and growth (see e.g. (7,11–13)). Cellular migration is a fundamental property of every cell and it is crucial for the development and morphogenesis of animal body plans and organ systems (see e.g. (14–16)). Cell migration is either in a random direction or directed towards localized cues (17–20). Mechanisms of cellular movement have been mostly studied in chemotactic cells, such as neutrophils (17), bacteria (21), Ciliata (22), fungi (23) and cellular slime molds (19).

Due to its extremely fast migration rate and highly irregular shape, the acellular slime mold *Physarum Polycephalum* represents a prime example of differentiated growth and thus offers an attractive model for the analysis of morphogenesis dynamics underlying cellular migration and exploration (24–28). *P polycephalum* is a giant single-celled organism that can grow to cover several square meters. Its morphology includes search fronts that are connected to a system of intersecting veins, in which oscillatory flows of the protoplasm “shuttle streaming” take place. This vein network allows 1) an efficient distribution of chemical signals, oxygen, nutrients over large distances and 2) cell migration at a speed of few centimeters per hour (29,30). The driving force for this protoplasm streaming is a periodic, peristaltic contraction and relaxation of the veins due to the actin-myosin interaction, which is regulated by oscillations of intracellular chemicals such as calcium (31–33). As it explores its environment, the slime mold extends temporary arm-like projections named pseudopods. It also secretes continuously a thick extracellular slime (34). The glycoprotein nature of the extracellular slime coat endows *P polycephalum* with unique protective and structural properties that favor survival of the migrating, naked slime mold (35). As the slime mold is foraging, it avoids areas covered with this mucus, which marks previously explored areas (36,37).

In the presence of chemical substances in the environment, *P. Polycephalum* shows directional movements towards or away from the stimuli (i.e. chemotaxis). *Physarum* morphology, evolution and behaviors are strongly affected by the availability, location and concentration of nutrients. When the slime mold senses attractants (e.g. food cues) using specific receptors located on the membrane, the oscillation frequency in the pseudopod closest to the attractant increases, causing cytoplasm to flow towards the attractant (38). On the contrary, when repellents such as salts are sensed, the oscillation frequency decreases and the slime mold moves away from the repellent (38). Although slime molds lack the complex hardware of animals with brains, they live in environments that are as complex and they face the same decision-making challenges (39). Hence, acellular slime molds have been the subject of a wide range of studies showing that they can solve complex biological and computational problems without any specialized nervous tissue (24,36,37,40–45).

In this paper the objectives are to characterize the morphology and dynamics of *Physarum* exploring various environments. First, we investigate how movement is affected by homogeneous environmental conditions: adverse environment (using salt as a repellent (46); nutritive environments (using glucose as a chemo-attractant (47,48) with 2 different concentrations) and a neutral environment (using plain agar). Second, we analyze the geometrical evolution of slime molds placed at a distance from a nutritive spot (glucose), with and without a repelling spot (salt) in between. We characterize slime molds’ movement both temporally and spatially, to capture the full dynamics. To this aim, we develop a program that automatically analyzes sequences of images to track the areas covered and explored by the slime mold, the slime mold shape, the refinement and secondary growth cycles, as well as the distance to the nutritive spot.

## Results

### 1) Homogeneous environment

In order to study the influence of the environment on slime mold expansion rate, we analyzed the areas covered by slime mold, unexplored substrate and mucus over time, as shown in Fig 1.

**Fig 1.**
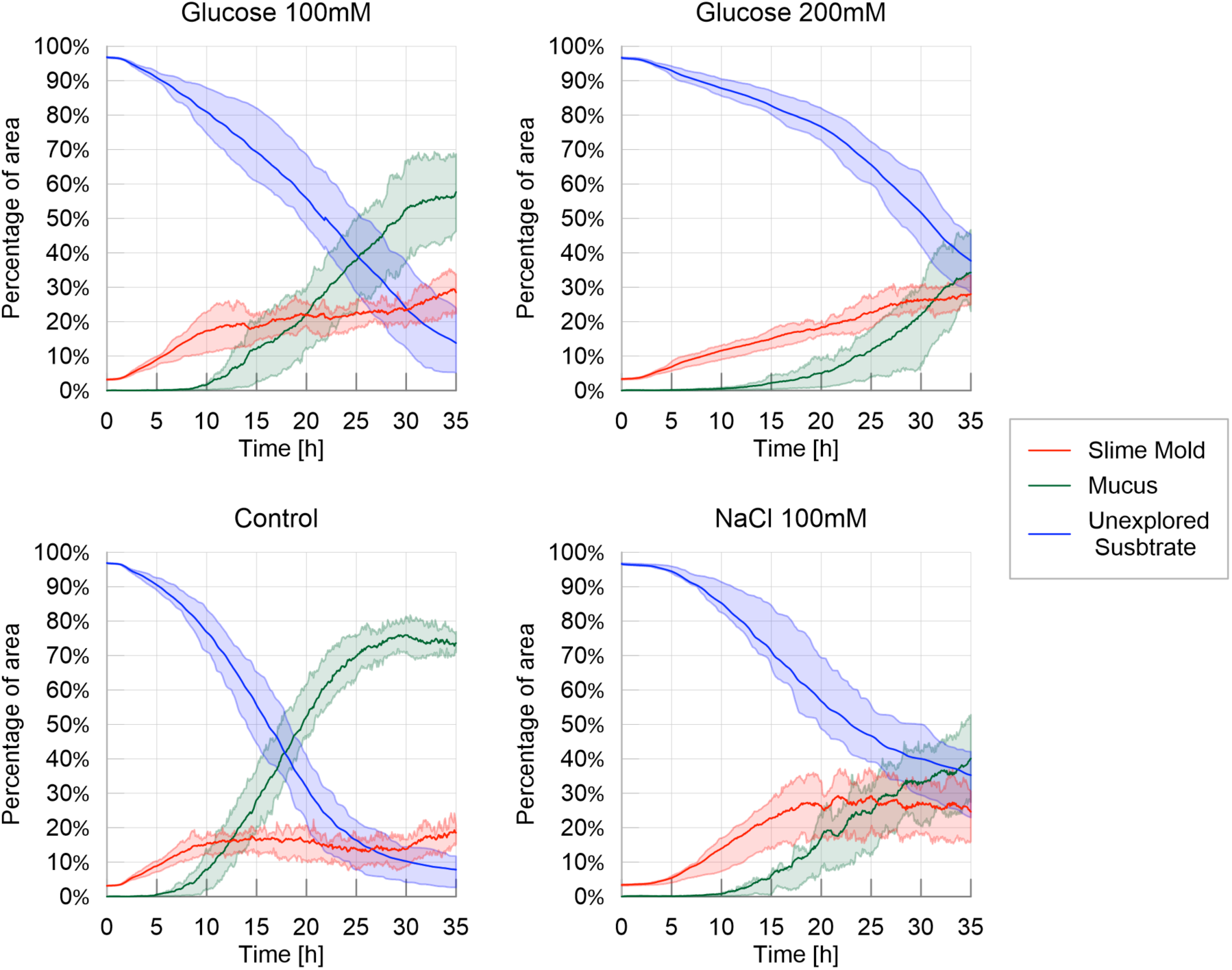
Fraction of area covered by slime mold, mucus and unexplored substrate – Homogeneous environment. The solid line corresponds to the average index calculated over the 20 replicates, while the shaded areas correspond to the first and third quartiles of the data.

In a neutral (control) and slightly nutritive environment (glucose at 100 mM), the slime molds started to spread from the very beginning of the experiment (Fig 1; Table 1 and Fig 5 in S1 Appendix, P>0.05). In contrast in a highly nutritive environment (200 mM glucose), slime molds started to explore later (P<0.001 when compared to the control). Slime molds placed in an adverse environment (100 mM NaCl) were lagging the most and only started exploring after 3 hours (P<0.001 when compared to the control). Once the slime molds started to explore, they all grew at the same rate (Fig 1; Table 2, Fig 6 in S1 Appendix, P> 0.05 when compared to the control) except the ones placed in a highly nutritive environment which were slowed down (P<0.001 when compared to the control).

At the end of the experiment (after 35 hours), the slime molds reached a similar surface in a control environment and in an adverse environment (Table 3 and Fig 7 in S1 Appendix, P>0.05 when compared to the control). Interestingly, after reaching a plateau at 18 hours, the area covered by the slime molds in an adverse environment oscillated with seemingly cyclic fluctuations (Fig 1). In both a slightly and a highly nutritive environment, the slime molds reached a higher final surface than the slime molds placed in a control environment (P<0.001 in both comparisons) and covered approximately 30% of arena at the end experiment. It is worth noting that in a highly nutritive environment, the surface covered by the slime molds never reached a plateau after 35 hours, suggesting that the slime molds did not reach its maximum surface (Fig 1).

Refinement *i.e.* appearance of mucus, was observed after 5 hours in the control environment. In all other environments, mucus appeared later (Table 4 and Fig 8 in S1 Appendix: P<0.001 for all treatments when compared to the control). In a highly nutritive environment, mucus was only observed after 10 hours, which marked the strongest delay in the refinement process. Once the mucus started to be apparent, its surface grew quicker in the control environment than in the other three treatments (Table 5 and Fig 9 in S1 Appendix; P<0.001 for all treatments when compared to the control). Thus the surface covered by mucus at the end of the experiment was the largest in the control environment where it reached 75% of the arena against 55%, 40% and 35% for the slightly nutritive, the adverse and the highly nutritive environments respectively (Table 6 and Fig 10 in S1 Appendix; P<0.001 for all treatments when compared to the control).

Hence, slime molds placed in a control environment explored almost all the arena leaving only 5% of the arena unexplored while in the other treatments the surface unexplored were significant: 15%, 35% and 38% for the slightly nutritive, the highly nutritive and the adverse environments respectively. Interestingly, although the growth rate dynamics differed between highly nutritive and adverse environments, the final unexplored surfaces were similar. In a highly nutritive environment the slime molds grew slowly and steadily while in an adverse environment slime molds grew rather quickly but after a long delay.

Next, we analyzed the evolution of the cumulative areas covered by primary growth, refinement and secondary growth (Fig 2). The cumulative area covered by secondary growth, which reveals the cyclic nature of the exploration process, was the highest in the adverse environment (480% coverage) followed by the control environment (380%), the slightly nutritive environment (250%) and the highly nutritive environment (180%). All comparisons lead to significant differences P<0.05, except control vs. adverse environment (Table 7 and Fig 11 in S1 Appendix). This observation confirms that exploration was slowed down by the presence of nutrients, and that the pulsatile behavior (*i.e* the exploration of previously explored area) was stimulated by repellents.

**Fig 2.**
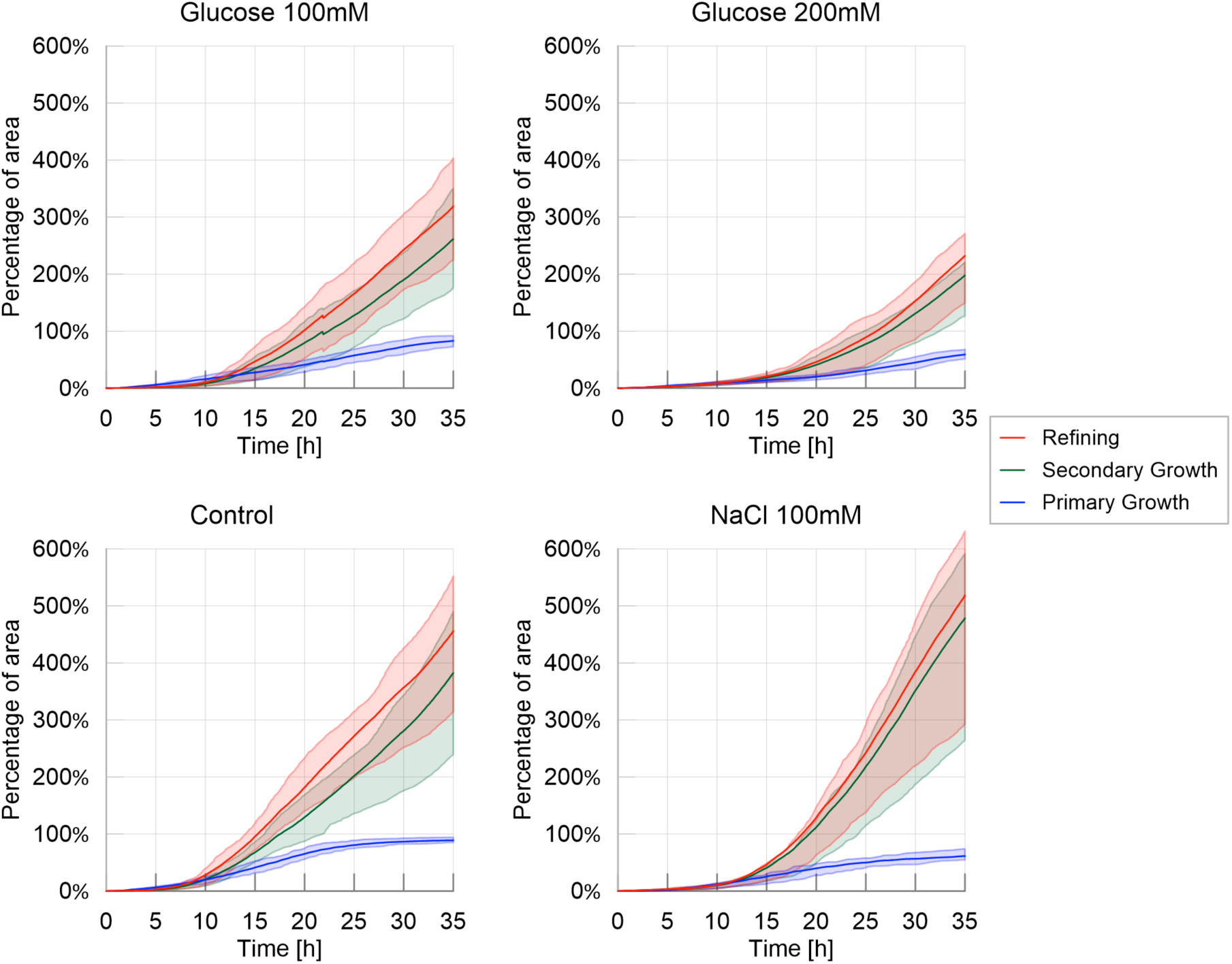
Cumulative areas covered by primary growth, refinement and secondary growth – Homogeneous experiments. The solid line corresponds to the average index calculated over the 20 replicates, while the shaded areas correspond to the first and third quartiles of the data.

**Fig 3:**
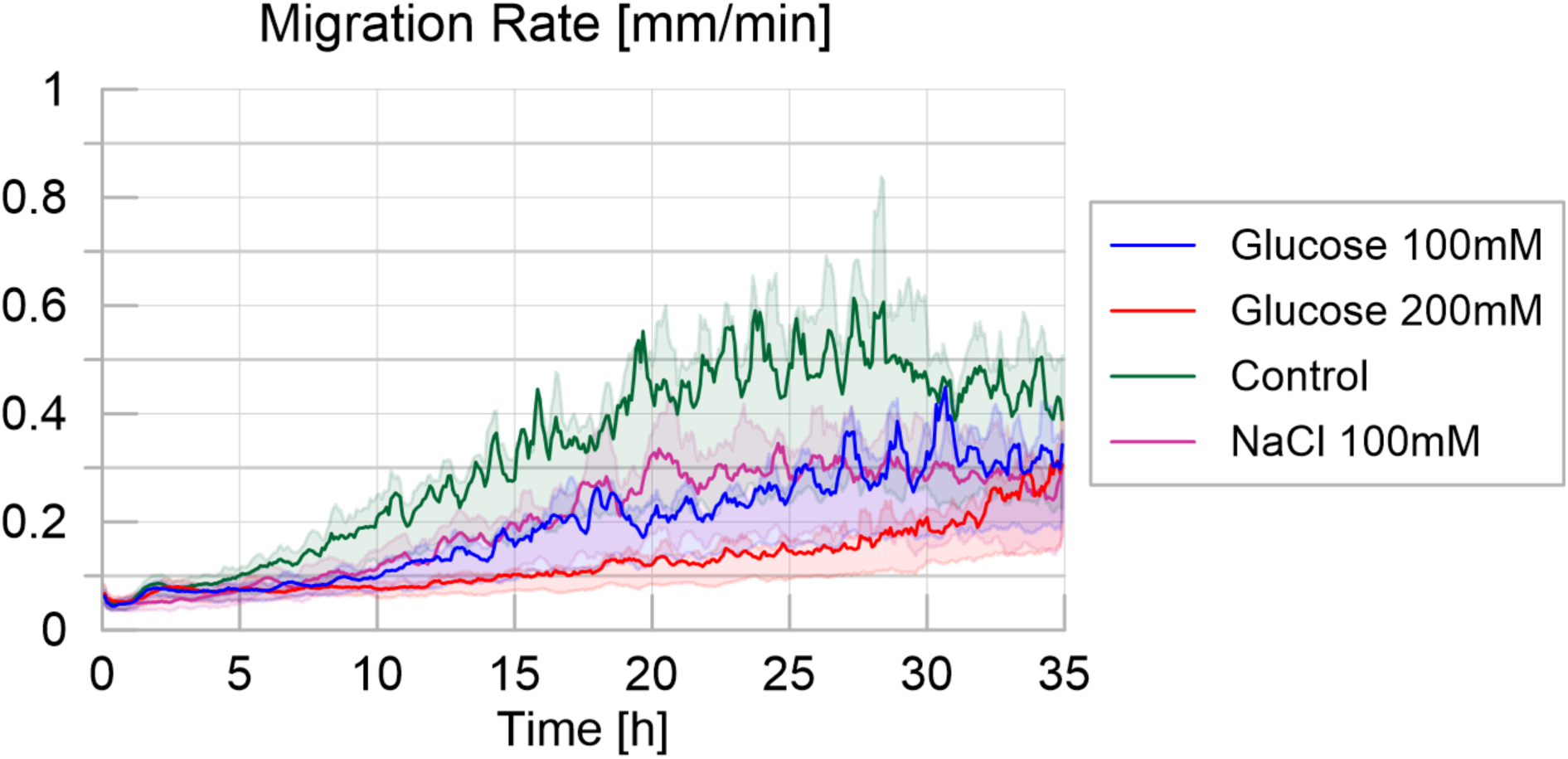
Migration rate over time for the four different treatments, defined as the maximum distance between the contours of the slime mold between two consecutive images divided by their time interval (5 minutes apart), measured in millimeters per minute. The solid line corresponds to the average calculated over 20 replicates per treatment, while the shaded areas correspond to the first and third quartiles of the data.

In accordance with the previous results, the migration rate was higher for the control treatment than for the other treatments (Table 8 and Fig 12 in S1 Appendix: P<0.001 for each pairwise comparison). While slime molds exploring the highly nutritive environment were slower than slime molds exploring the slightly nutritive or the adverse environment (P<0.001 each), these two showed no significant differences (P>0.05).

The slime molds exploring the adverse environment showed the highest probability to explore a previously explored substrate than the other treatments as shown in Fig 4 and supplementary materials (Tendency for secondary growth: Table 9 and Fig 13 in S1 Appendix: P<0.001 for each pairwise comparison). When exploring a highly nutritive environment, slime molds also displayed a significant positive tendency for secondary growth (P<0.001 for each pairwise comparison) but significantly less strong than on the adverse environment (P<0.001). For the others treatments, the measured proportion of secondary growth was not different from the expected proportion of secondary growth, indicating that the slime molds did not avoid previously explored substrate and explored randomly (Fig 13 in S1 Appendix). The peaks observed within the first 5 hours of the experiment correspond to an isotropic extension immediately followed by a refinement process that occurred before the slime mold started to explore continuously its environment. This behavior is often observed when a slime mold is introduced in a new environment and is referred as “contemplative” (49) *i.e.* the slime mold migrates, retracts and moves again. The peak was larger for adverse environment.

**Fig 4:**
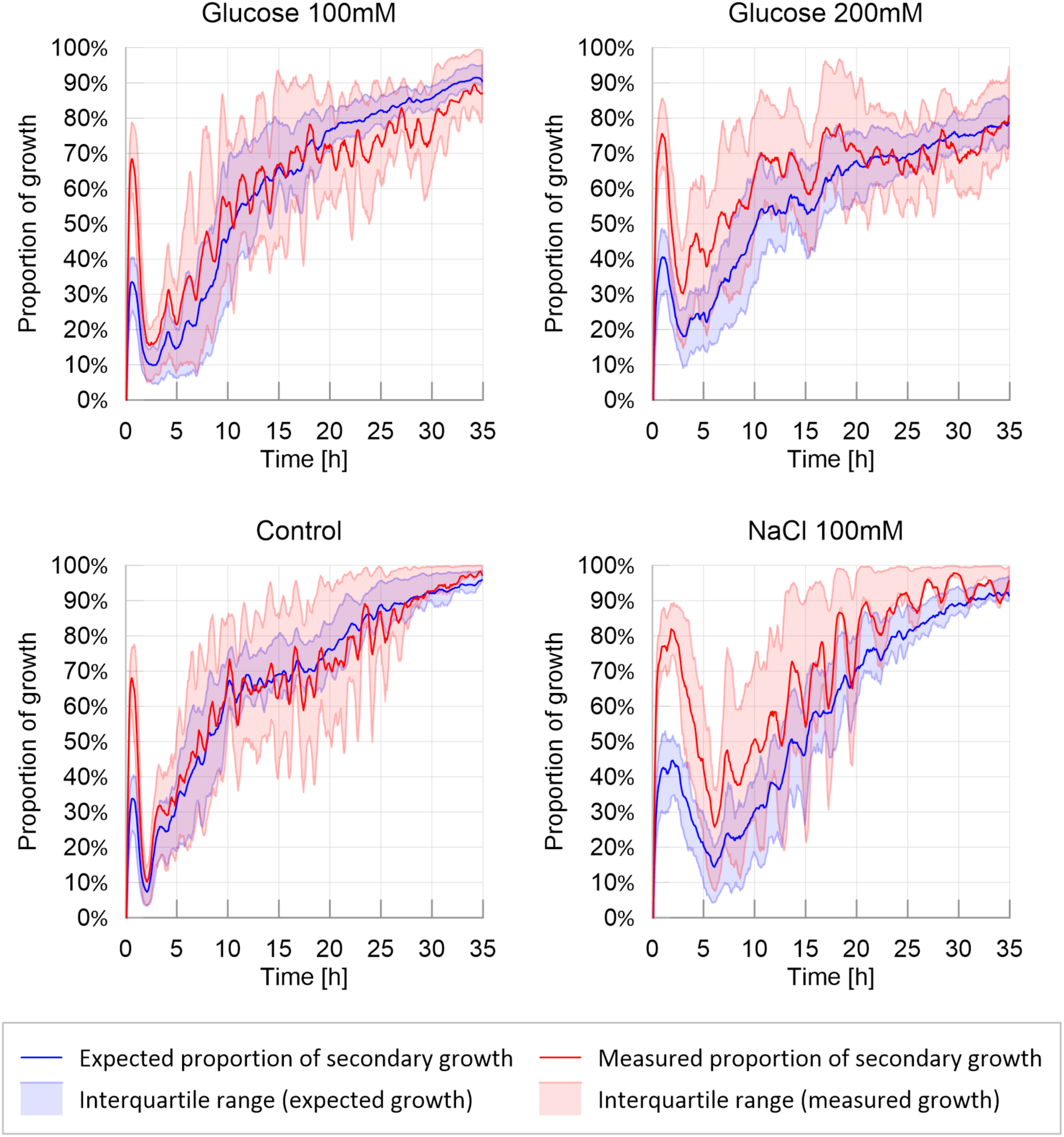
Ratio of secondary growth: observed and expected proportion of secondary growth. The solid line corresponds to the average calculated over 20 replicates per treatment, while the shaded areas correspond to the first and third quartiles of the data.

We then analyzed the evolution of the shape of the slime molds contour. Note that the experimental set ups in which slime molds were placed exhibited radial symmetry. Hence, no preferential expansion direction was expected. We thus focused on contour shape, not orientation.

As expected, circularity was initially one in all tests (circular slime mold spot), and increased over time as the contour shape departed from a circle (Fig 2 in S1 Appendix). In the control and nutritive environments, circularity remained between 1.05 and 1.10, whereas it fluctuated between 1.05 and 1.30 in the adverse environment. This observation suggests that, in an adverse environment, slime molds explored the petri dish by spreading and thinning over larger areas than in the other environments, which led to shape changes and a decrease of slime mold circularity. However fluctuations among the 20 replicates were too high to identify any trend in the evolution of slime mold circularity.

The eccentricity index was initially close to zero (circular cell) and increased up to almost 0.8 over time, with important fluctuations in all the treatments (Fig 3 in S1 Appendix). Eccentricity is an indicator of the number of pseudopodia. But a non-eccentric convex hull can enclose non-circular contours of slime mold, since pseudopodia can develop in a symmetric fashion. That is why no major difference was noted between the treatments. This result highlights the absence of preferred expansion direction in symmetric, homogeneous environments.

Solidity decreased with the emergence of pseudopodia, since slime mold branching disrupted the initially convex shape of the slime mold (Fig 5). In the control environment, in which the exploration rate was the highest, the decrease of solidity of the slime mold area was the highest (and the fastest) decreasing from 1 to 0.3 and then becoming relatively stable, with fluctuations of +/- 0.05 (Table 10 and Fig 14 in S1 Appendix; P<0.01 for all paired comparisons except adverse environment vs. slightly nutritive environment, where P>0.05). Slower exploration resulted in a slower and steadier loss of solidity as observed in the nutritive and adverse environments. The highly nutritive environment yielded the highest solidity at the end of the experiment (0.6), which confirmed that the presence of glucose slowed down exploration.

**Fig 5.**
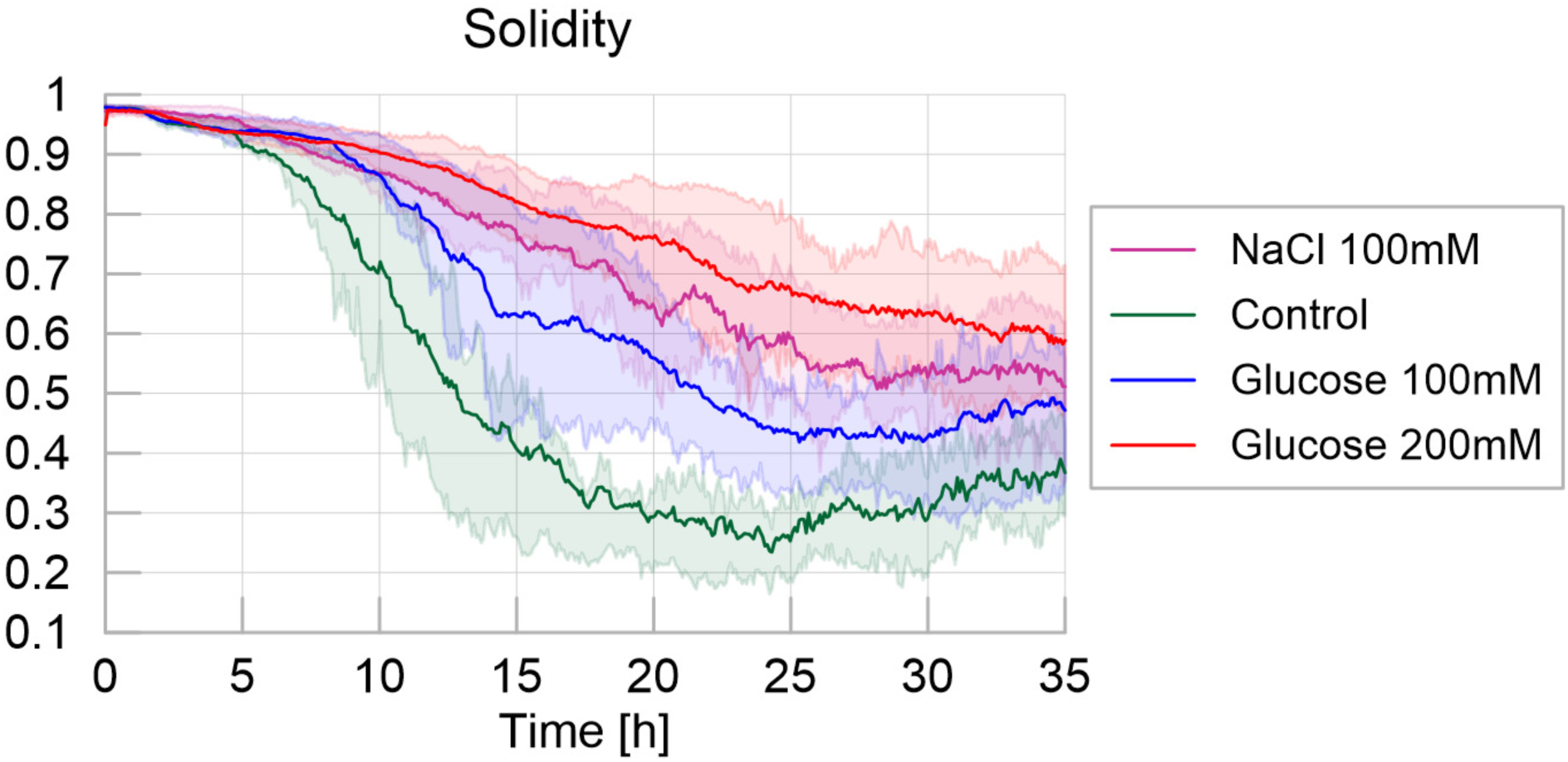
Solidity - Homogeneous experiments. The solid line corresponds to the average index calculated over the 20 replicates, while the shaded areas correspond to the first and third quartiles of the data.

We next focused on the number of clusters, corresponding to the number of pseudopodia (Fig 6). Initially the slime molds stretched as a single cluster. Once mucus started to be apparent, slime molds usually divided up into several clusters, and started the active exploration phase. In highly nutritive and adverse environments, the number of clusters over time was lower than in the other two treatments (new pseudopod number: Table 11 and Fig 15 in S1 Appendix, and P < 0.01 for all paired comparisons except for adverse environment vs. highly nutritive environment). This observation confirmed that the presence of concentrated nutrients slowed down the exploration, and that the presence of repellents triggered a highly pulsatile behavior with small exploration fronts, which were sometimes not detected as separate clusters.

**Fig 6.**
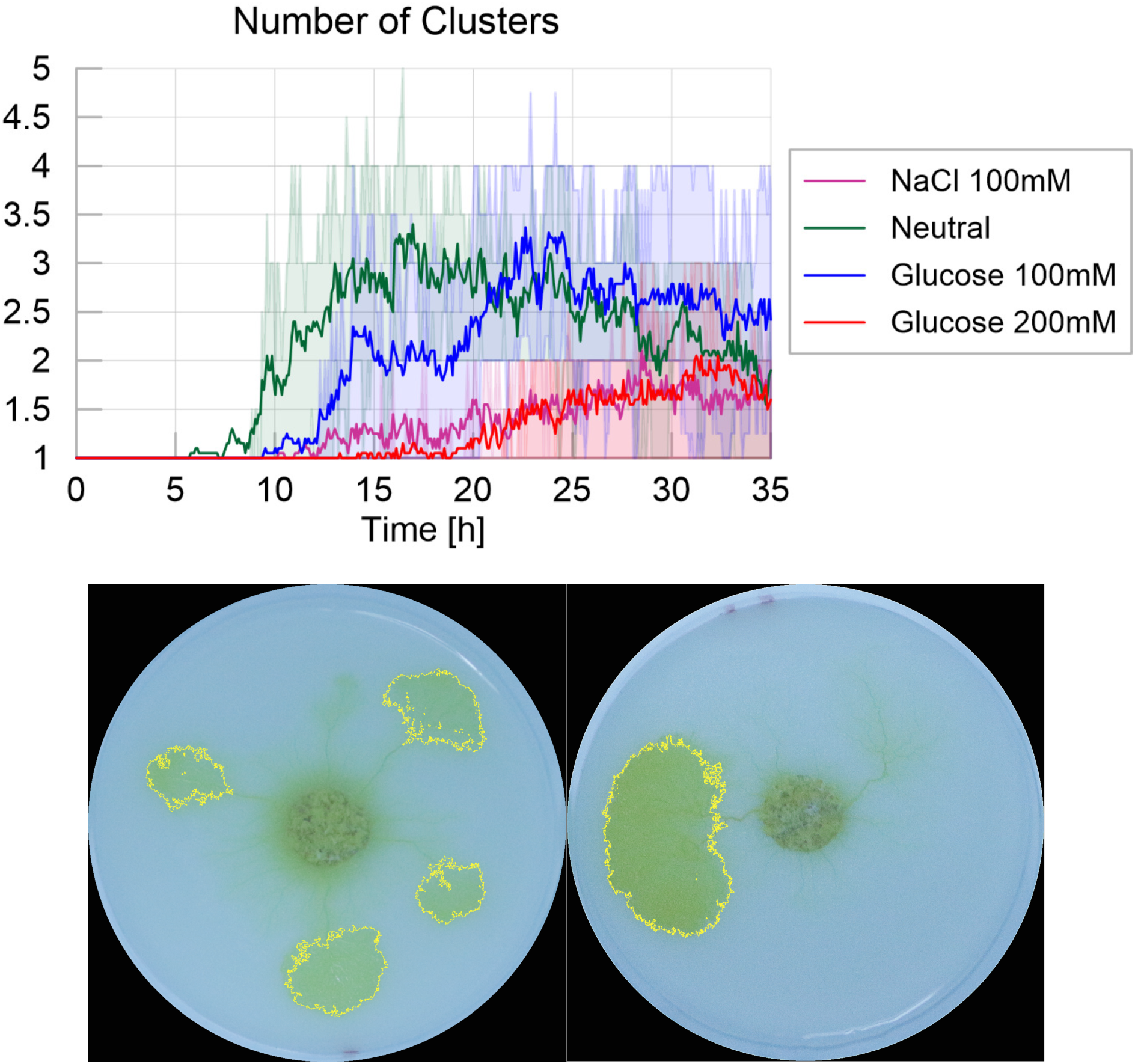
Number of clusters - Homogeneous experiments. The solid line corresponds to the average index calculated over the 20 replicates, while the shaded areas correspond to the first and third quartiles of the data. Pictures show examples of clusters for the Control environment (left) and 200 mM Glucose environment (right).

### 2) Spot experiments

In the spot experiments, we studied the influence of discrete distributions of nutrients and repellents on exploration dynamics. When looking at the evolution of slime mold, mucus and unexplored substrate over time (Fig 7), we only observed marginal difference among the treatments, which all exhibited similar patterns of exploration, e.g. similar percentage of non-explored area and similar mucus accumulation. The presence of an adverse spot only delayed the appearance of the first pseudopod (first movement: Table 12 and Fig 15 in S1 Appendix, P<0.05) but not the first appearance of mucus (first appearance of mucus: Table 15 in S1 Appendix, not significant). The only noticeable differences lie in the surface reached at the end of the experiment: slime molds that were offered a highly nutritive spot grew larger (final surface: Table 14 and Fig 17 in S1 Appendix; P < 0.01). By contrast, the surface covered by mucus was lower (mucus final surface: Table 17 and Fig 18 in S1 Appendix; P <0.01) than slime molds that were offered a slightly nutritive spot. In comparison with the experiments conducted in homogeneous environments, we did not observe any expansion/refinement cycles in the spot experiments, meaning that slime mold spread steadily towards the food source despite the presence of an obstacle on the way.

**Fig 7.**
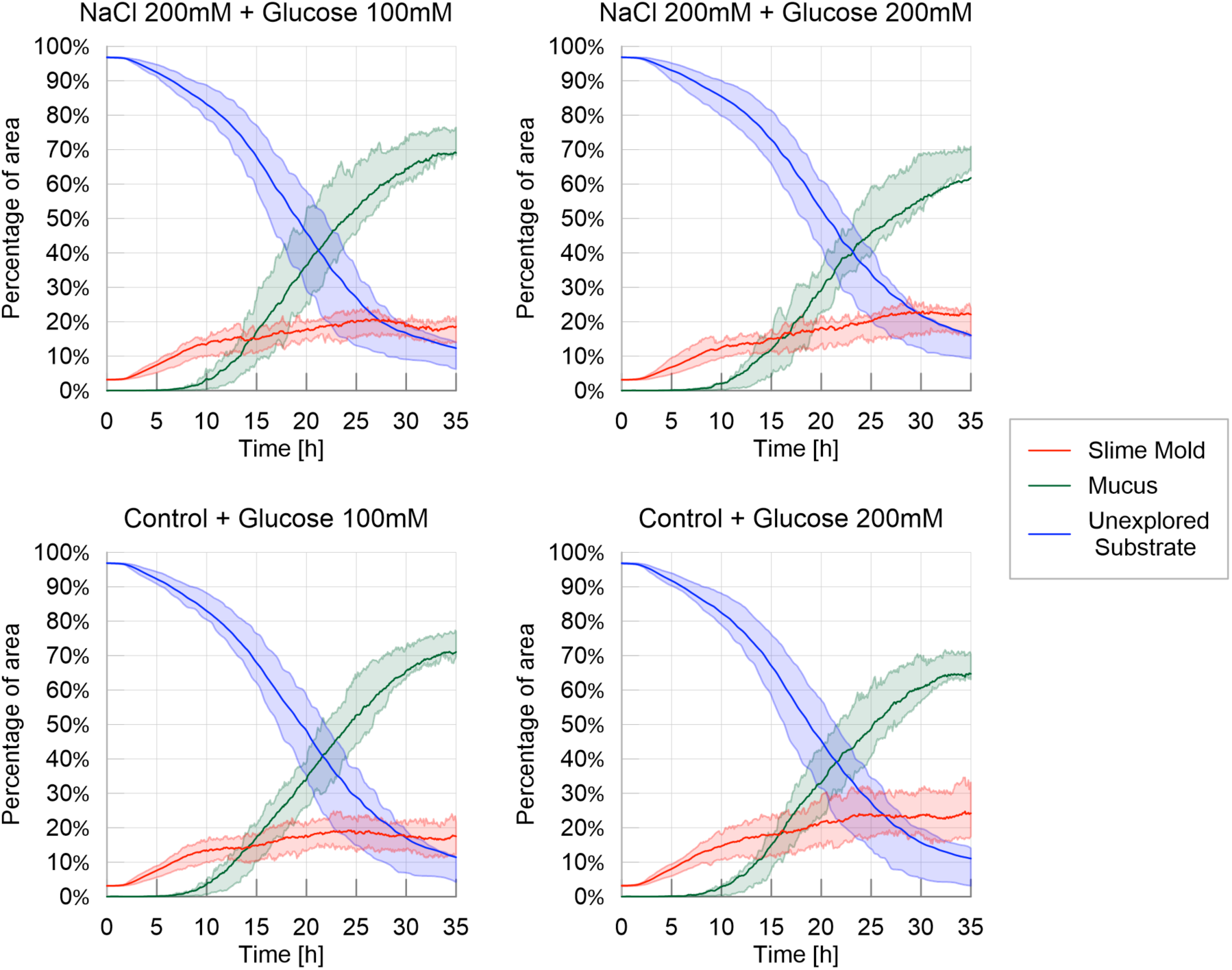
Fraction of area covered by each entity (slime mold, mucus, unexplored substrate) – Spot experiments. The solid line corresponds to the average index calculated over the 20 replicates, while the shaded areas correspond to the first and third quartiles of the data

The exploration behavior in the spot experiments was similar to that observed in the control environment in homogeneous experiments as observed in Fig 8, which includes the average percentage of unexplored area for the homogeneous and spot experiments shown in Fig 1 and Fig 7 respectively. This suggests that the spatial exploration of slime mold depended mostly on the substrate and not on the geometric distribution of the nutritive and adverse stimuli.

**Fig 8.**
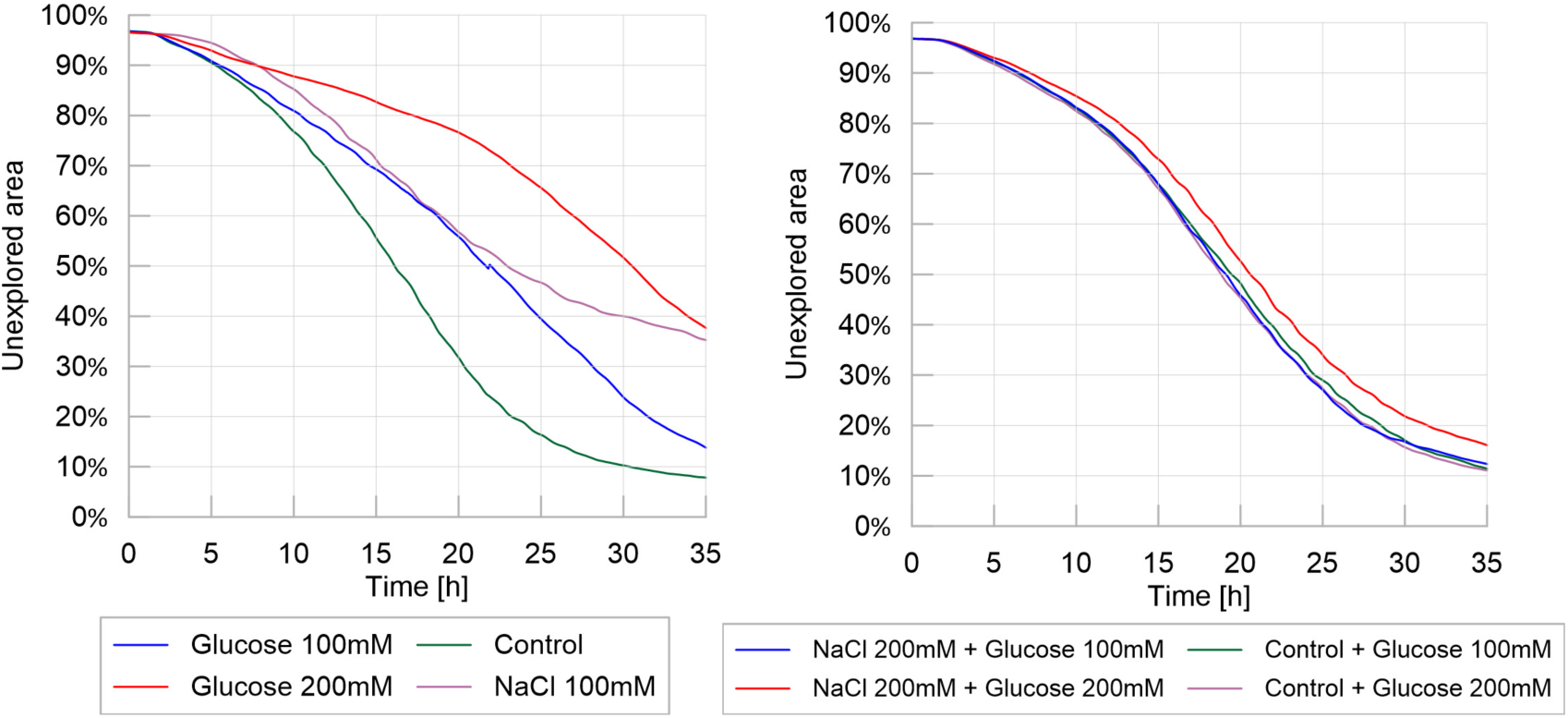
Exploration behavior: homogeneous experiments (left) vs. spot experiments (right). Percentage of unexplored area over time. Mean values of over 20 replicates for each different treatment.

The cumulative areas covered by secondary growth for the spot experiments (Fig 9) were also similar for all treatments (Table 18 in S1 Appendix, P >0.05), suggesting again, that isolated spots with nutritive or adverse stimuli did not alter the overall exploration of slime molds when growing on the same, control, substrate.

**Fig 9.**
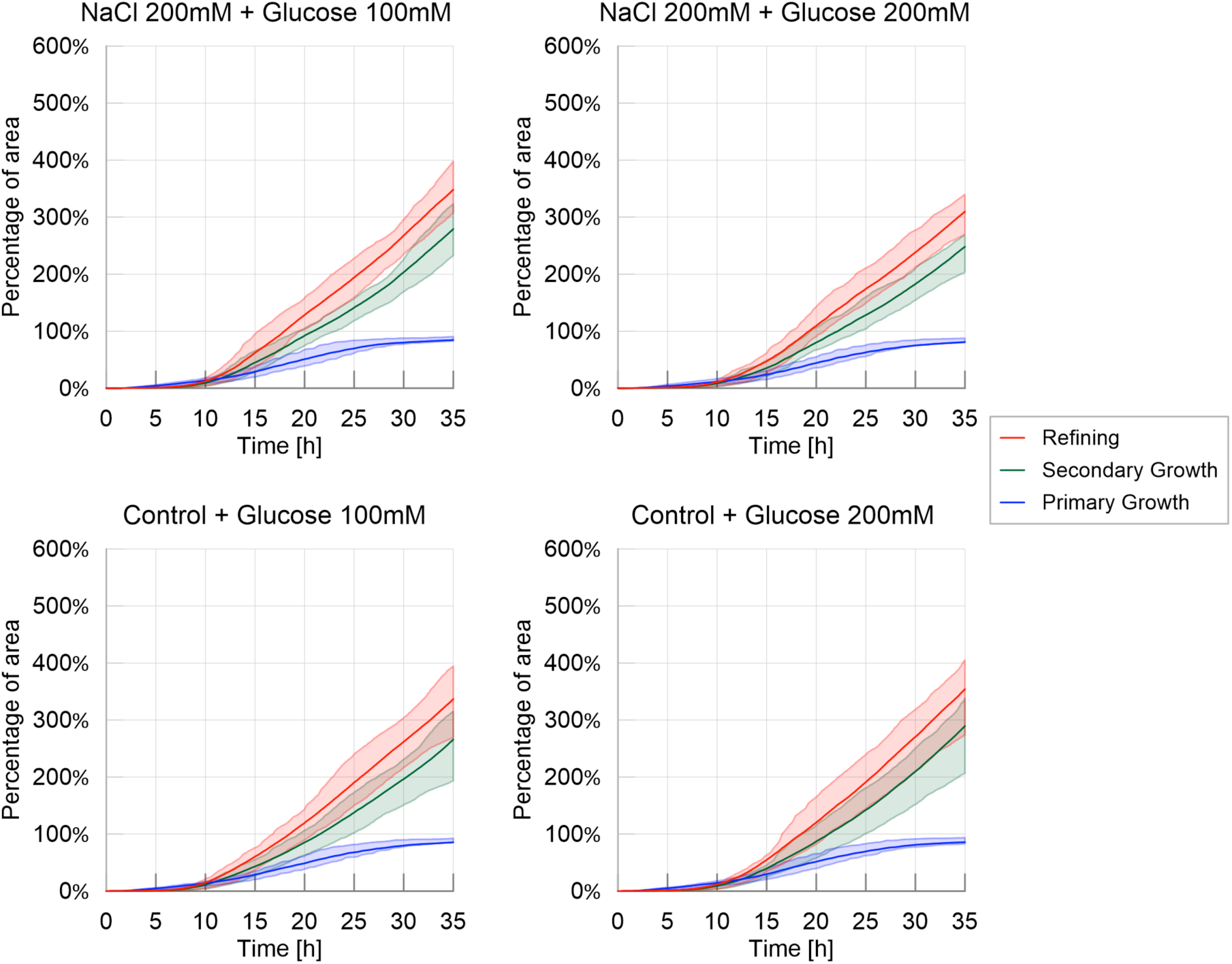
Cumulative areas covered by primary growth, refinement and secondary growth – Spot experiments. The solid line corresponds to the average index calculated over the 20 replicates, while the shaded areas correspond to the first and third quartiles of the data

The trends of migration rate, as shown in Fig 10, show that slime molds were not affected by the difference between treatments, showing only a slight effect of the concentration of the food spot (P<0.05), as shown on Table 19 in S1 Appendix. This effect showed that the migration rate was slightly superior when slime molds were offered a higher than a lower nutritive spot.

**Fig 10:**
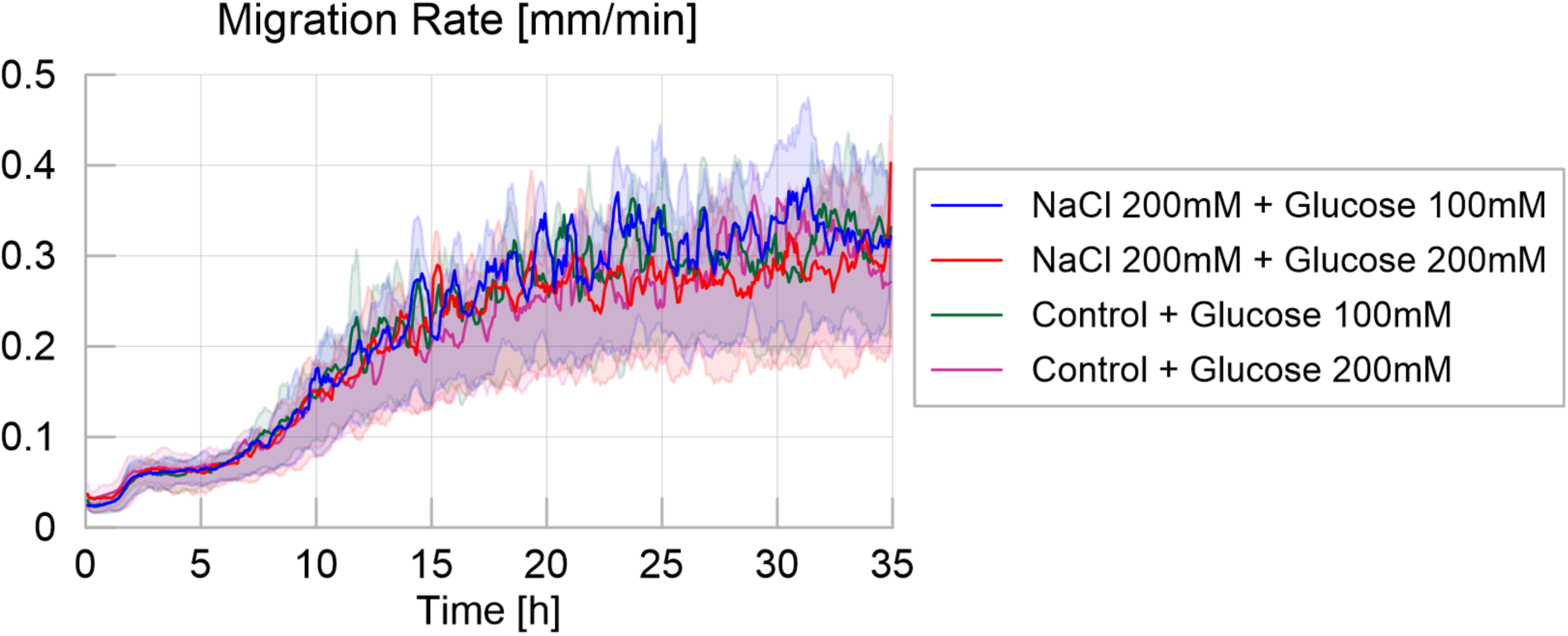
Migration rate over time for the four different treatments, defined as the maximum distance between the contours of the slime mold between two consecutive images, divided by their time interval (5 minutes apart), measured in millimeters per minute. The solid line corresponds to the average calculated over 20 replicates per treatment, while the shaded areas correspond to the first and third quartiles of the data.

Similarly, looking at the predilection of slime molds to grow towards mucus (secondary growth), as shown in Fig 11, no significant differences were observed between treatments (Table 19 in S1 Appendix), which suggests that the growth type is not influenced by the existence or concentration of discrete attracting/repelling spots.

**Fig 11:**
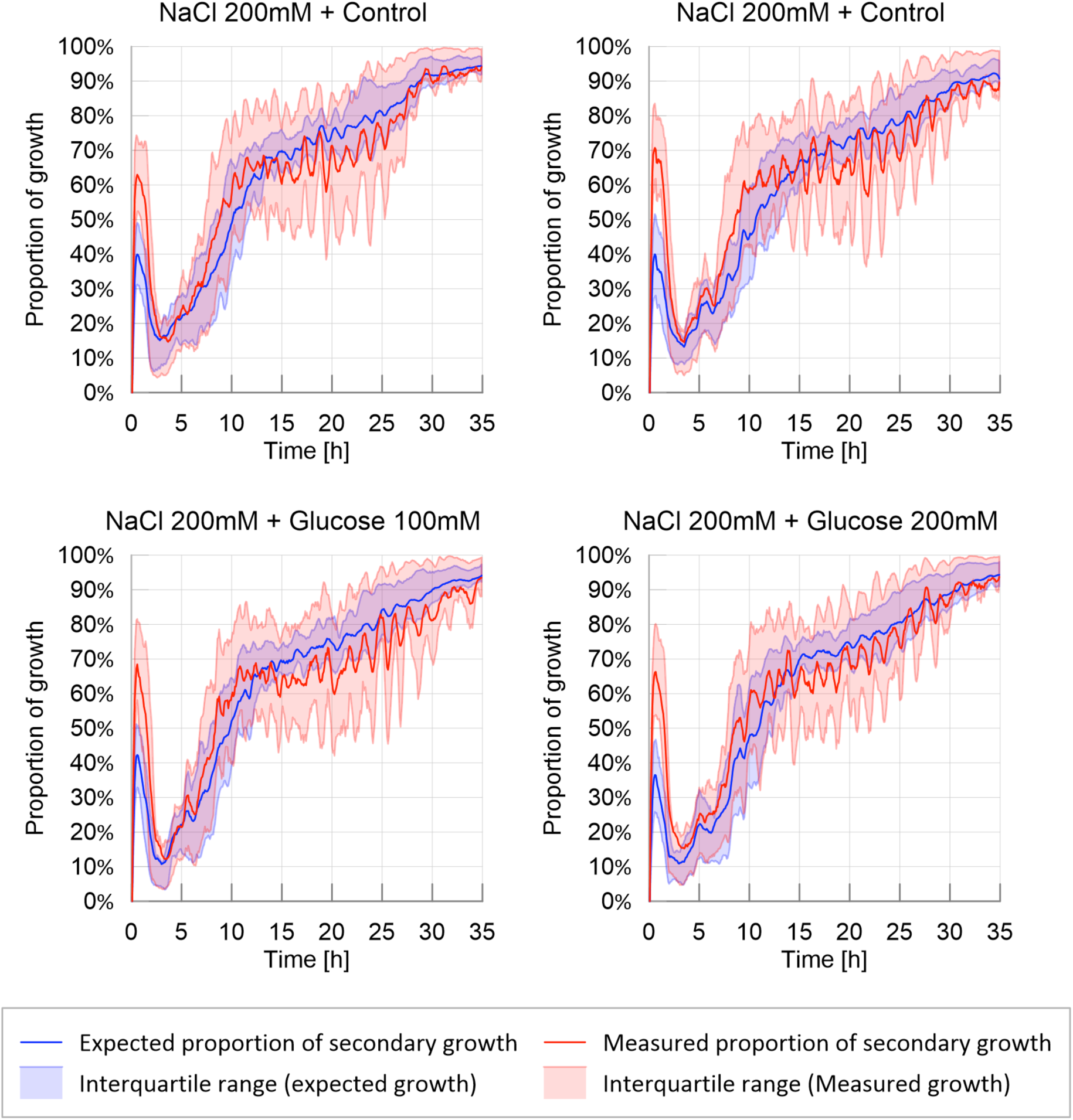
Probability of secondary growth: observed and expected proportion of secondary growth. The solid line corresponds to the average calculated over 20 replicates per treatment, while the shaded areas correspond to the first and third quartiles of the data.

The results obtained for the four different shape indexes (Fig 4 in S1 Appendix) for the spot experiments support the hypothesis that discrete spots of nutrients or repellents did not affect the overall expansion dynamics and exploration cycles. This interpretation is confirmed by the average number of pseudopods, which was found to be correlated to the formation of mucus during the exploration phase: in all the spot experiments, the number of clusters increases from 1 to 2.5 within around 12 hours, to reach a plateau afterwards. In other words, less exploration cycles were observed in non homogenous environments, yielding less pseudopodia.

The evolution of the shortest distance from the slime mold cell to the glucose spot is shown in Fig 12, which can be viewed as a “survival” plot, displaying the proportion of the replicate (P) in which slime mold has not reached the nutritive spot at a given time. Both increasing the concentration of nutrient in the spot (from slightly nutritive to highly nutritive) and adding an aversive spot increased the time to reach the food patch by decreasing the probability to reach it (time to food patch: Table 22 and Fig 18 in S1 Appendix, nutritive: P< 0.01; aversive: P< 0.05).

**Fig 12.**
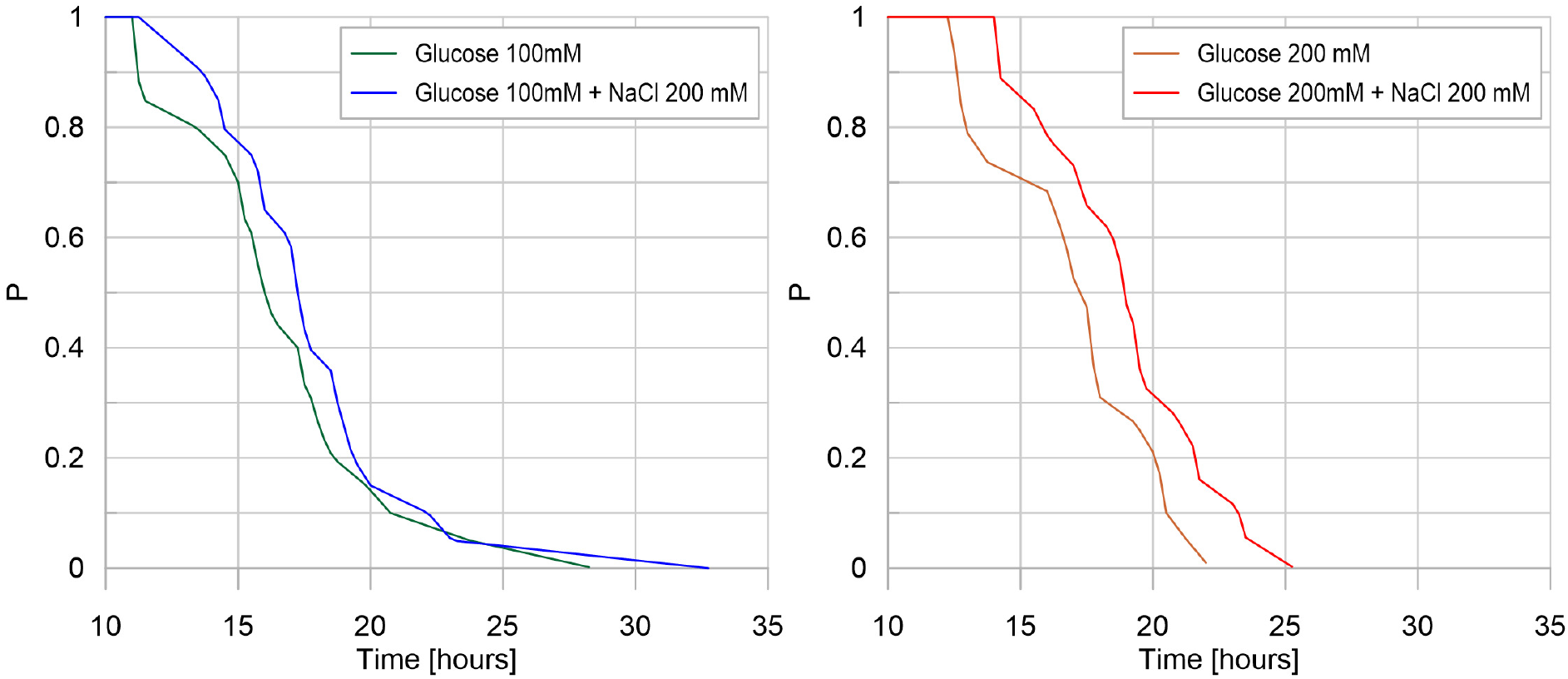
Survival plot: glucose concentration effect. For each treatment, on the vertical axis, the value P corresponds to the fraction of replicates that have not reached the glucose spot at a given time. For a representative number of replicates, P is the probability that glucose has not been reached by slime mold at a given time, for a specific treatment. On the horizontal axis, each value of time corresponds to the average time it takes for a certain fraction of slime mold (P) to reach the glucose spot.

## Discussion

Exploration in slime molds involves two different processes: area extension and movement (27). Slime molds locomotion and morphogenesis depend on the response of the organism to the environmental conditions. In the present paper, we show that distributions of nutritive and aversive cues affected drastically the exploration pattern of slime molds.

The typical exploration behavior of a slime mold (control condition) started by a stretching period where the slime molds grew uniformly in all directions for 10 hours. Then, the contour of slime mold lost circularity when the first pseudopodia appeared, which also corresponds to the first occurrence of mucus. This phenomenon is typical of the directed digitated growth, or branching phase, described by numerous authors (27,51,59–61). Slime molds developed multiple pseudopodia and did not exhibit any preferential exploration orientation. At the end of the experiment, almost all the arena was explored by the slime mold. Thus the exploration was characterized by three phases: (i) Primary growth only, in the quasi-absence of mucus (5-10 hours); (ii) Combination of primary and secondary growth; (iii) Secondary growth only, when the slime mold stops exploring new substrate areas.

### 1) Homogeneous distribution of nutritive cues

First, we noted that an environment containing a uniform distribution of nutrients slowed down the exploration of slime mold, mainly by delaying secondary growth and increasing the period of the pulsatile exploration/refinement movements. The area not explored by the slime molds was 3 times larger (respectively 7 times larger) than in the environment deprived of nutrient (control case) for a slightly nutritive (respectively highly nutritive) environment. The exploration rate was almost linear for highly nutritive environments, while for other treatments, the area covered by the slime molds reached a plateau after a period of stretching, which indicates secondary growth and slime mold displacement. This means that slime molds that explored nutritive environments never exhibited a Phase (iii) in their exploration pattern.

Second, on substrates with higher nutrient concentrations, the slime molds grew in a more compact fashion, *i.e*. slime molds presented the highest solidity index and the lowest number of pseudopodia (clusters). Additionally, the appearance of mucus, which indicates that the slime mold was withdrawing, occurred much later in nutritive environments. As glucose is only aversive when only above 300 mM (54), our results suggest that nutritive media depressed migration due to feeding behavior. This allows the organism to remain at a site until nutrients are exhausted (54,62,63). In previous studies, it was shown that the area of substrate covered increases when slime mold responds to nutrient dilution (54,64). Here, we confirmed these observations and noted that slime mold tended to migrate and grow faster on substrates with the lowest concentration of nutrients, thus maximizing nutrient intake and optimizing the trade-off between nutrient foraging and nutrient intake.

### 2) Homogeneous distribution of aversive cues

The aversion of slime mold to salt manifested itself through longer contemplative behavior, delayed primary growth and a higher probability to crawl on previously explored surface. In addition, the slime molds grew more compactly and with less pseudopods. This suggests that slime molds were actively avoiding contact with the aversive surface (44,49). In the absence of cell walls, slime mold has no other protection from the environment than mucus. Indeed, in bacteria for instance, mucus is used as protective barrier for the cells against harsh external conditions (65). In slime molds, the extracellular mucus excreted by the slime molds can have different roles: hydrophilic shield to prevent water loss (66), a lubricating surface over which the slime molds can easily crawl (67), a defensive coat to protect against invasion by foreign materials and organisms (66,68), an aid to phagocytosis (69), a surface that promotes ion-exchange (66) and has externalized spatial memory that helps navigation in unknown environments (26,36,53). Here, we can add a new function for the mucus i.e. a buffer to move in adverse environment.

### 3) Non-Homogeneous distribution of aversive and non aversive cues

In the spot experiments, pulsatile movements were limited and slime molds responded in the same way regardless of the concentration of glucose used as attractant and regardless of the presence of a salt spot on the way to the glucose spot. Slime molds grew in a more compact fashion, *i.e*. slime molds presented the highest solidity index and the lowest number of pseudopodia (clusters). Additionally, the evolution of the areas covered by slime mold and mucus over time indicates that the response of slime molds to heterogeneous environments was similar to that in the control case. This result suggests that in the spot experiments, the exploration behavior of slime mold is mostly controlled by the substrate. Our observation confirms that salt reception can be affected by the presence of sugars (46). The authors in (46) showed that the “apparent” enthalpy change accompanying salt perception decreases with increase of sugar concentration.

The proposed image analysis program allows extracting information on expansion rates, geometric changes and probability of occupancy. Ongoing developments aim to acquire high quality images of slime mold exploration tests and to expand the code’s capabilities to extract topological information on the networks formed by slime molds. Possible applications of the code go beyond the study of slime molds. For instance, the analysis of successive images of an ant nest could allow detecting the generation of any rhythmic activity. Trajectories of individuals within a flock could also be described to understand whether or not members of the flock can inform and influence the travel direction of other individuals. Such a finding would allow understanding how group decisions are made among gregarious species. The cluster identification and shape recognition program could be used to differentiate modes of gene expression or to recognize objects. Object identification is an important pillar to explain associative memory or to track species interactions in an ecosystem. The tools and approach presented here are thus applicable to any problem of network dynamics or pattern recognition.

## Methods

### 1) Species

*Physarum Polycephalum*, also known as the true slime mold, belongs to the Amoebozoa, the sister group to fungi and animals (50). Slime molds are found on organic substrates like tree bark or forest soil where they feed on microorganisms such as bacteria or fungi (50). The vegetative morph of *P. polycephalum*, the plasmodium, is a vast multinucleate cell that can grow to cover up to a few square meters and crawl at speeds from 0.1 to few centimeters per hour (29,30). When hygrometry and food availability decrease, the plasmodium turns into an encysted resting stage made of desiccated spherules called sclerotium (29).

### 2) Rearing conditions

Experiments were initiated with a total of 10 sclerotia per strain (Southern Biological, Victoria, Australia). We cultivated slime molds on a 1% agar medium with rolled oat flakes, slime molds were fed every day and the medium was replaced daily. Slime molds were 2 weeks old when the experiment started. All experiments were carried out in the dark at 25°C temperature and 70% humidity, and ran for 35 h. Pictures were taken every 5 min with a Canon 70D digital camera.

### 3) Experimental setup

Initially we monitored the exploration movement evoked in slime molds in a homogeneous environment. Each slime mold was placed in the center of a circular arena (14.5 cm in diameter) with a layer of agar (1% in water) mixed with non-nutritive cellulose (5%). Adding cellulose to the agar mix proved to be useful to obtain a homogeneous pigmentation and to enhance the color contrast between the substrate and slime mold, therefore improving the identification process. A circular hole (2.5 cm in diameter) was punched and replaced with a circular slime mold of the same size sitting on oat. In the first and second treatments (nutritive environments) we added glucose (100 mM or 200 mM) to the medium. In the third treatment (adverse environment), we added a known repellent (NaCl 100 mM (51)) to the medium. Lastly, in the fourth treatment, the medium remained unchanged (neutral environment *i.e.* control treatment).

Subsequently, to investigate how chemotaxis modified the exploration behavior, we introduced discrete spots of attractants/repellants within a neutral substrate made of plain agar. In these so-called “spot experiments”, we followed a procedure similar to that for the homogeneous environments. A circular slime mold (2.5 cm in diameter) was placed diametrically opposite to a glucose (attractant) spot of same size placed 4.5 cm away. In some of the treatments, a salt (NaCl 200 mM, repellent) spot was added at the center of the petri dish. A total of 4 different treatments were tested: the first and the second with a single spot of glucose at concentrations of 100 mM and 200 mM respectively, the third and fourth keeping the glucose spots with the same concentrations and adding a NaCl 200 mM spot at the center of the dish.

All slime molds were fed just before the experiment so we assumed that they were in the same physiological state. The experiment consisted of a total of 8 different treatments. We replicated the experiment 20 times for each treatment and monitored each arena for 35 hours taking time-lapse photographs every 5 minutes. Fig 13 shows the experimental set-up for homogeneous environments (left) and discrete distributions of attractants/repellants (right).

**Fig 13:**
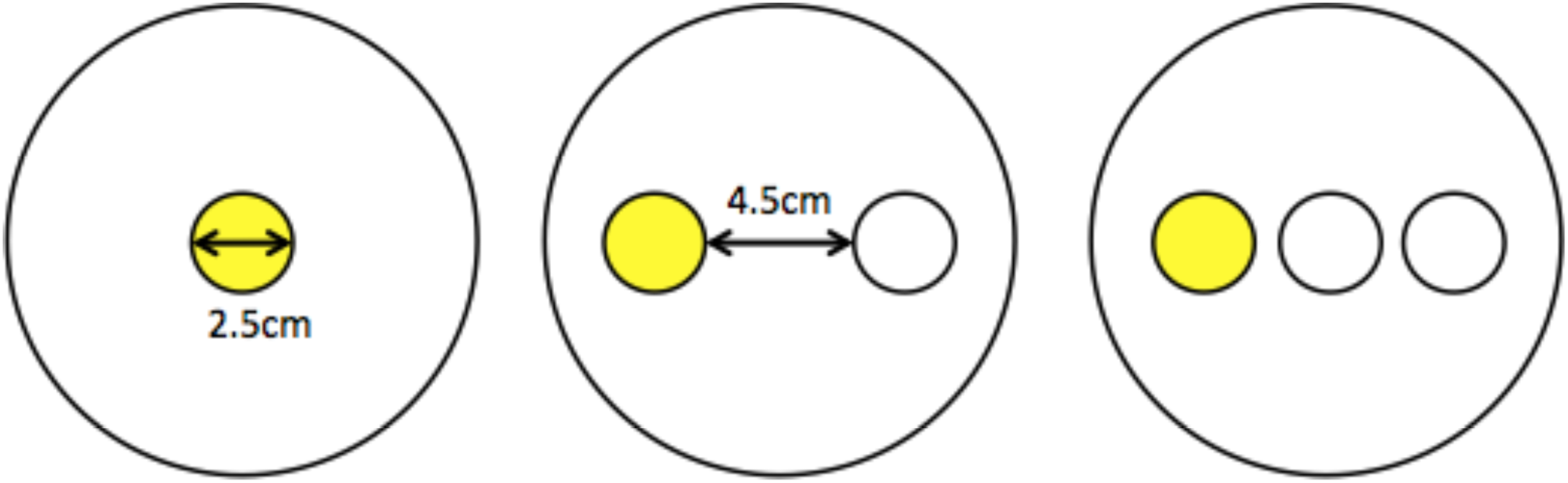
Experimental set-ups for homogenous environment and spot experiment

### 4) Image Processing

Time-lapse images were taken every 5 minutes for a total of 420 pictures for each replicate. The images acquired initially belong to the RGB color space and their size was 1200 by 1200 pixels. Image analysis followed three main steps. First, the edge of the petri dish was identified by fitting its border to a circle of known diameter. Second, the image was segmented using the clustering algorithm k-means (52), which was applied to the images converted into the ab* color space, which is the CLAB space without the L* (lighting) component; this choice corresponds to the robustness of this color space against changes of lighting conditions between images. By the end of this step, all the pixels inside petri dish, e.g. the region of interest (ROI), are identified as either slime mold or not-slime mold.

After distinguishing slime mold and non-slime mold areas, we identified the mucus by using a subtraction method. The mucus is left by slime after refinement; this substance acts as a marker present on already explored areas of the domain, serving as an external memory to the slime mold (36,53). The fact that this mucus is transparent makes it very challenging to identify by sole color analysis. We thus trinarized the image based on the history of a given pixel, since a pixel that is classified as non-slime-mold at the current image will necessarily contain mucus if it has ever been classified as slime mold in any of the previous images. Conversely, a non-slime-mold pixel will be classified as unexplored substrate if it has never hosted slime mold up to that point in time.

The change of class from unexplored substrate to slime mold, defined as primary growth, means that a new sprout of the slime mold reaches a point in space that it had never explored before. Similarly, secondary growth is defined as the change from mucus to slime mold, meaning that the slime mold is revisiting an already explored location. Lastly, if the slime mold recedes from a point, e.g. a pixel goes from slime mold to non-slime mold, it becomes mucus, and the process is defined as refinement. By the end of these three steps, the images have been trinarized, meaning that every pixel is classified as slime mold, unexplored substrate, or mucus (and the points outside the ROI are disregarded). Two videos are provided to the reader as supplementary material, S2 and S3 show one of the replicates as original images and trinarized images after identification respectively.

### 5) Image Analysis in Space and Time

After completing the trinarization, we calculated several indexes to characterize slime mold geometry dynamics. We averaged the indexes over the 20 replicates of each treatment in order to obtain statistically representative measures, and we plotted them over time.

In order to quantify the differences of slime mold spreading dynamics on distinct substrates, we first calculated the fraction of the petri dish area covered by slime mold, mucus and unexplored substrate over time. The total area, the lighting conditions and the test duration were the same for all treatments, both in the homogeneous and spot experiments. Note that glucose only provides energy to slime mold, which is not gaining significant mass during the experiments (54). In other words, slime mold is changing its area by mostly by stretching and contracting, therefore changing its area density.

In order to gain further insights about the exploration process we then computed the cumulative area of primary growth, refinement, and secondary growth over the full period of the experiments comparing two consecutive images at the time. The cumulative area covered by primary growth is indicative of the total area of exploration, therefore it is always smaller or equal to the total area of the dish. The cumulative area covered by secondary growth indicates whether slime mold expansion is monotonic (dominated by primary growth) which results in a smaller magnitude, or cyclic (secondary growth dominated, with pulsatile movements) which results in a larger magnitude. The cumulative area covered by refinement indicates slime mold density changes. Within a given time interval, if the area covered by primary plus secondary growth equals that covered by refinement, then slime mold keeps the same density, whereas if it is superior, the slime mold stretches (e.g. density decreases). If secondary growth is negligible and if the area covered by primary growth equals the area covered by refinement, then slime mold displaces mass.

We next measured the migration rate. To this aim, for two consecutive images, we measured the distance from each pixel where growth occurred (both primary and secondary) to the closest pixel classified as slime mold in the previous image. This distance represents the extent of growth from one image to the next. We calculated the migration rate as the ratio between the maximum distance traveled and the time interval between two images (5 min). This maximum distance traveled was then used to delineate the region explored by slime mold within the 5 min interval. In other words, we defined an area of interest as the contour of the slime mold with an offset corresponding to the maximum distance traveled (see Fig 1 in S1 Appendix for more details).

We estimated the fraction of secondary growth as the ratio between the number of pixels changing from mucus to slime mold and the total number of pixels in the region of interest. We then calculated the fraction area of “expected secondary growth”, which would have occurred if secondary growth had happened randomly. If the measured secondary growth fraction is higher (respectively, lower) than the expected one, this means that slime mold has a bias towards mucus (respectively, unexplored substrate).

Additionally, we computed four shape parameters indicative of the contour of slime mold: circularity, eccentricity, solidity and number of clusters. Circularity (C) is defined as:

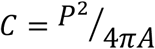

Where P and A are the perimeter and area of the shape of slime mold at a given time; this index is equal to one when the contour of slime mold is circular, and increases as the shape deviates from the circle. Eccentricity (E) is calculated as the ratio between the distance between the foci and the major axis length, as follows:

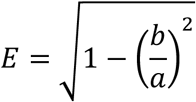

In which *a* and *b* are the lengths of the major and minor axes, respectively. When E is equal to zero, the contour is a circle; when E is equal to one, the contour is degenerated into a line. Solidity (S) is the ratio between the area of the slime mold contour and the area of its convex hull, e.g. the smallest convex polygon that encloses all the slime mold pixels. S is equal to unity when the contour shape is convex. Lastly, we measured the number of clusters by performing an “erosion” operation along the contour of the slime mold, consisting in removing the veins that connect the regions of high concentration of slime mold. After this erosion process, only the clusters of high concentration of slime mold remained, which provided the number of pseudopodia at the given image.

For the spot experiments, we also determined the distance from the slime mold to the glucose spot at every time step. This distance was calculated as the minimum distance between the contour of the slime mold and the perimeter of the glucose circular spot. The evolution of the distance to glucose over time was analyzed in a way similar to a survival analysis, as described below.

### 5) Statistics

The full description of the statistics is provided as part of the supplementary information; Appendix S1 includes the results of the complete statistical analyses, while appendix S4 gives the necessary instructions to reproduce those analyses. When dependent variables lasted until the occurrence of certain event, we conducted survival analyses using the R package coxme (55). For the remaining dependent variables, we did linear analyses using the R packages lme4 (56) and lmerTest (57). For the experiments done in homogeneous environment, we tested the four different treatments (control, nutritive and adverse) as fixed factors. For the spot experiments, we tested the effect of the nature of the nearest spot (neutral or NaCl 200 mM) and that of the furthest spot (glucose 100 mM or 200 mM), as well as the interaction between the two. We always considered the date of the experiment as a random factor as all treatments were conducted over multiple days, with several replicates per day for each treatment. For each statistical analysis, we performed a nested model comparison using the R package MuMIn (58) by ordering models according to their Akaike criterion. We represented the selected model by plotting estimators with the pairwise comparison (a posteriori Tuckey test) p-values using the R package emmeans (58) in linear models, and the hazard ratio associated p-values in cox models.

## Supporting information

**S1 Appendix. Supplementary information: Image analysis and statistical results.** Description of the methodology to extract indexes from image analysis and results obtained from the statistical analysis that support our observations.

**S2 Video. Time-lapse of one experiment replicate, original images.** Video showing the evolution of the slime mold cell over the 35 hours of the experiments, original acquired photos.

**S3 Video. Time-lapse of one experiment replicate, trinarized images.** Time lapse of the results of the image segmentation for the same experiment shown in S2 Video.

**S4 Appendix. Statistic analysis script description.** Step by step description of the procedure followed to generate the results of the shown statistical analyses.

## Acknowledgements

This work was supported by the U.S National Science Foundation, under grant CMMI#1552368: “CAREER: Multiphysics Damage and Healing of Rocks for Performance Enhancement of Geo-Storage Systems - A Bottom-Up Research and Education Approach.” A.D. and A.B. were supported by a grant from the Agence Nationale de la Recherche (reference number: ANR-17-CE02-0019-01-SMART-CELL).

